# Tethered agonist activated ADGRF1 structure reveals molecular preference for Gα_q_ signalling

**DOI:** 10.1101/2022.09.09.507336

**Authors:** Daniel T. D. Jones, Andrew N. Dates, Shaun D. Rawson, Maggie M. Burruss, Colin H. Lipper, Stephen C. Blacklow

## Abstract

Adhesion G-Protein Coupled Receptors (aGPCRs) have evolved an activation mechanism to translate extracellular force into liberation of a tethered agonist (TA) to modulate cell signalling. We report here that ADGRF1 is the first class B GPCR shown to signal through all major G-protein classes and identify the structural basis for its Gα_q_ preference by cryo-EM. Our structure shows that Gα_q_ over Gα_s_ preference in ADGRF1 derives from tighter packing at the conserved F569 of the TA, altering contacts between TM helix I and VII, with a concurrent rearrangement of TM helices VII and VIII at the site of Gα recruitment. Gα_s_ signalling is also more sensitive to mutation of TA or binding site residues than Gα_q_. Our work advances the understanding of aGPCR TA activation in molecular detail, identifying structural features that potentially explain preferential signal modulation.

## Main

Adhesion G protein coupled receptors (aGPCRs) are the second largest class of GPCRs, constituting 33 members across 9 subfamilies ^1^. aGPCRs control a multitude of cellular processes involved in organ development and tissue homeostasis, responding to external stimuli that sense extracellular physical forces, such as binding to protein ligands presented on cells, components of extracellular matrix and shear flow^2^. Mutations in aGPCRs are genetically responsible for human developmental disorders, such as ADGRV1 in Usher syndrome^3^, and EMR2 in vibratory urticaria^4^.

aGPCRs have a unique modular architecture, with an N-terminal ectodomain containing extracellular adhesive modules, followed by a G-protein coupled receptor autoinducing (GAIN) domain linked to a seven-transmembrane (7TM) bundle that relays extracellular events to the cell. A distinguishing hallmark of aGPCRs is that the GAIN domain undergoes autoproteolysis at a short hairpin turn between the final two beta-strands, separating the aGPCR into non-covalently attached N-terminal and C-terminal fragments, named NTF and CTF respectively^5,6^. Currently, models for aGPCR activation suggest that mechanical force applied to the adhesive modules in aGPCRs induces dissociation of the NTF from the CTF^7,8^, enabling the newly liberated N-terminal end of the CTF to act as an intramolecular agonist to the 7TM domain. Intramolecular ligation of this tethered agonist (TA) to the 7TM domain relays the initial extracellular cue to the cell through G-protein coupling or beta-arrestin activity.

The first structure of an aGPCR 7TM domain was solved for ADGRG3 (GPR97) bound to glucocorticoids in the orthosteric pocket, revealing key residues involved in ligand recognition, conformational switches important for receptor activation, and residues for coupling to G-protein^9^. Cryo-EM structures were then reported for seven different aGPCR family members activated by their TAs, including representative GPCR-G-protein complexes for all major families of Gα, except for Gα_q_ ^10–13^. These studies showed that the TA acts an intramolecular ligand for the 7TM domain and identified a canonical binding pose for a conserved hydrophobic interaction motif (HIM) of the TA in the orthosteric site.

ADGRF1 (GPR110) is an aGPCR belonging to aGPCR subfamily VI ^1,14^. ADGRF1 was identified as an oncogene overexpressed in lung and prostate cancer^15^, and has been implicated in synaptamide dependent neural outgrowth and repair^16,17^, though this activity remains controversial^18^. ADGRF1 was used as a prototypical aGPCR to determine that membranes expressing ADGRF1 treated with urea at high concentration induced shedding of the NTF, liberating the TA to activate the 7TM and recruit Gα_q_ preferentially over other Gα proteins^8^. Further work has also showed that mouse ADGRF1 can be activated by synthetic TA peptides when added in trans, signalling through both Gα_s_ and Gα_q_^19^.

Recent structures of the TA-engaged state of the ADGRF1 CTF coupled to either miniGα_s_ and miniGαi also identified a lysophosphatidylcholine (LPC) lipid bound at the intracellular convergence of TM helices II, III and IV^13^. Interestingly, fragments of ADGRF1 expressed with the GAIN domain included also yielded TA-engaged structures of ADGRF1 coupled to miniG but lacked density for the GAIN domain, suggesting that the CTF may have dissociated from the NTF portion of the protein during purification.

To understand the molecular events driving TA activation of ADGRF1 and how its structure determines Gα class preference, we determined a cryo-EM structure of TA-activated ADGRF1 coupled to miniGα_s/q_-β_1_γ_2_, stabilised by Nb35. Combining structural insights with extensive functional studies, we identify key residues of the orthosteric binding site that are essential for Gα_s_ signalling but dispensable for Gα_q_ signalling. Overall, our work complements other recent aGPCR structures and extends them by visualizing ADGRF1 in complex with its most relevant G-protein partner, elucidating a functional and structural landscape by which ADGRF1 is activated and uncovering how its structure determines Gα class preference. Furthermore, the mechanistic insights determined here regarding TA engagement to strong and weakly associated G-proteins offers valuable insight into how TA mediated activation of aGPCRs may be functionally modulated.

## Results

### ADGRF1 signals through all G-protein classes

We recently developed a platform for analysis of the signalling activity of aGPCRs, in which the CTF of interest is expressed as a protein fusion with an N-terminal IL2 signal sequence followed by maltose binding protein (MBP) and a tobacco etch virus cleavage site (denoted MBP-CTF) ^20^ (Extended Data Fig. 1). Using this approach to measure signalling activity in reporter assays, we observed transcriptional responses with both CRE and NFAT reporters, indicative respectively of Gα_s_ and Gα_q_ signalling, consistent with previous studies^13,21^. We also observed transcriptional responses with the SRE and SRF-RE reporters, which suggests that ADGRF1 can also signal through Gα_i/o_ and Gα_12/13_ (Fig. 1a).

**Figure 1.**
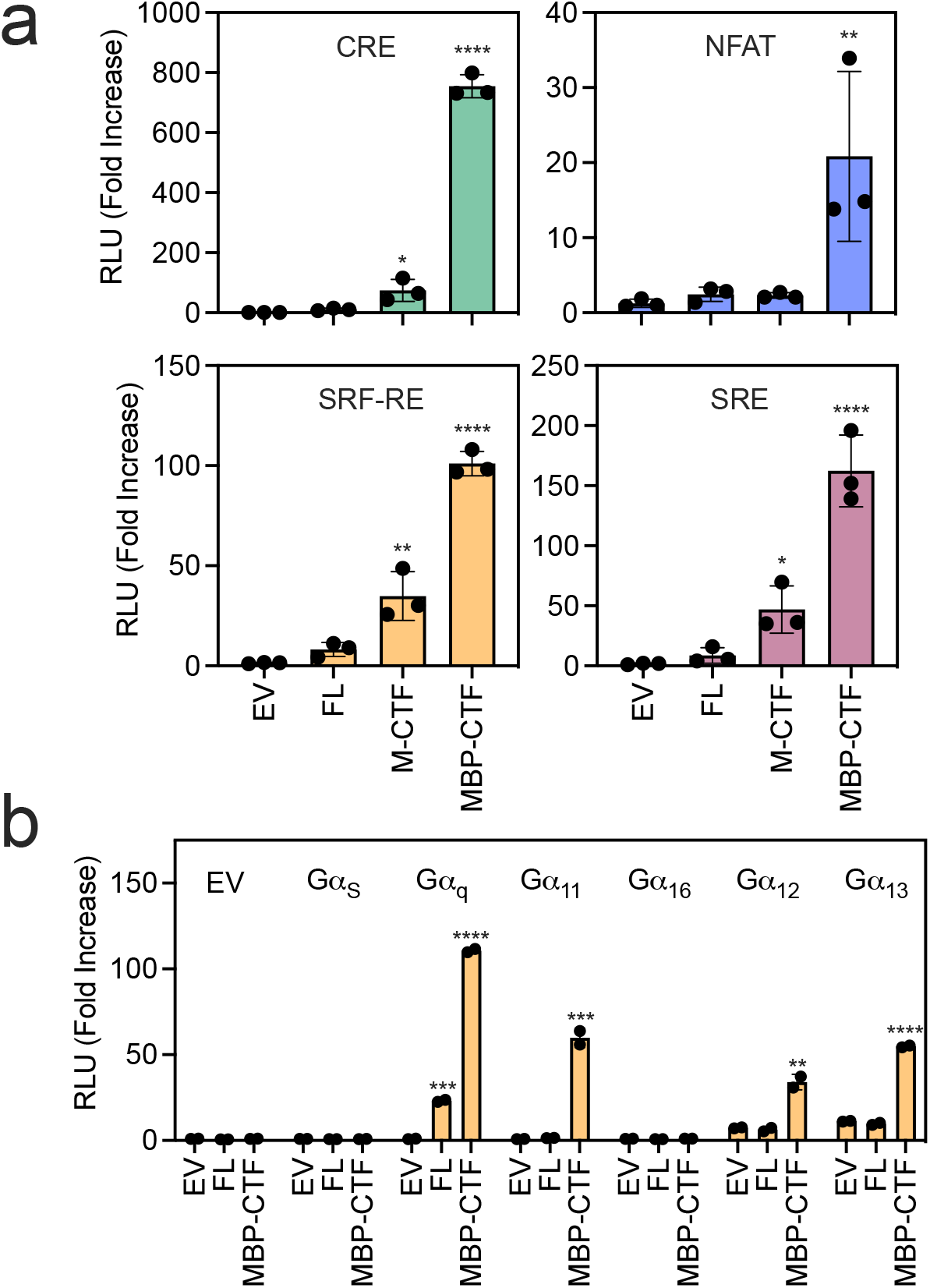
ADGRF1 signals through all G-protein classes. (a) Transcription factor reporter assay responses for ADGRF1 FL, M-CTF or MBP-CTF relative to empty vector for CRE (top left), NFAT (top right), SRF-RE (bottom left) and SRE (bottom right). Cells were transfected with 30 ng DNA/well for SRE and SRF-RE, and 10 ng DNA/well for CRE and NFAT. Data are normalized to empty vector as relative luminescence units (RLU). Data are presented as mean ± s.d. of three biological replicates. One-way analysis of variance (ANOVA) was used with Dunnett’s multiple-comparison post-hoc test to compare the difference between samples to each reporter’s empty vector control (*P≤0.05, **P≤0.01, and ****P≤0.0001). (b) Gα transient transfection complementation assay. Activity of the SRF-RE reporter was measured in HEKΔ6 cells transfected with a combination of ADGRF1 and different Gα subunits. The Gα subunit or Gα empty vector control was co-transfected with receptor empty vector, ADGRF1 FL or MBP-CTF. Values are reported as the ratio of the raw luminescence values of firefly and renilla luciferases. Data are presented as mean ± s.d. of two biological replicates. Each point represents the mean value of three technical replicates. In all panels, One-way analysis of variance (ANOVA) was used with Dunnett’s multiple-comparison post-hoc test to compare the difference between samples to each Gα empty vector control (*P≤0.05, **P≤0.01, ***P≤0.001 and ****P≤0.0001).

To further evaluate the potential for signalling through Gα_12/13_ (a structure of Gα_i_ has been reported^13^) we used HEK293T cells with deletion of six Gα subunits (Gα_s_ short, Gα_s_ long, Gα_q_, Gα_11_, Gα16, Gα_12_ and Gα_13_, gift from Associate Prof. A. Inoue, Tohoku university, Japan)^22^, and deletion complementation by transient transfection to determine which Gα subunits support ADGRF1 signalling in the SRF-RE reporter assay. ADGRF1 MBP-CTF signals strongly through Gα_q_ and Gα_11_, and also exhibits a smaller significant signal for Gα_12_ and Gα_13_ (Fig. 1b). These data show that ADGRF1 has the functional capacity to signal through all Gα classes, making ADGRF1 the first of the class B GPCRs shown to have this capability^23^.

### Protein engineering of ADGRF1 and Cryo-EM structure determination

To obtain an active state ADGRF1 G-protein protein complex amenable for structural and functional investigation, we probed complex assembly with a suite of miniGα proteins^24,25^, using a NanoBiT recruitment assay^26, 27^. We found that LgBiT-miniGα_s_, miniGα_s/q_ and miniGα_o_ were all recruited to C-terminal SmBiT-tagged ADGRF1-CTF as judged by increased luminescence upon SmBiT-LgBiT complementation, whereas LgBiT-miniGα_12_, and matched CΔ5 controls were not (Extended Data Fig. 2a). To assess tractibility for structure determination, we co-expressed these constructs in Expi293T suspension cells, and also observed a similar pattern of Gα recruitment to ADGRF1 in these cells (Extended Data Fig. 2b, left). To further test the stability of the complex, we solubilised cells with mild detergent supplemented with cholesterol in the presence of apyrase.

The only complex that retained luminescence under these lysis conditions was the combination of SmBiT-tagged ADGRF1-CTF with LgBiT-miniGα_s/q_ (Extended Data Fig. 2b, right). In addition, only miniGα_s/q_ specifically purified with ADGRF1-CTF using anti-FLAG M2 magnetic beads (Extended Data Fig. 2c), indicating that ADGRF1-Gα complex stability is greater for miniGα_s/q_ compared to miniGα_s_ and miniGα_o_. Furthermore, miniGα_s/q_ co-expression with ADGRF1-CTF appears to increase the amount of receptor purified, suggesting that the miniGα_s/q_ may improve receptor expression by facilitating the folding of the receptor, increasing its stability, or by blocking toxic signalling through endogenous signalling pathways.

### Structure of the ADGRF1-miniGα_s/q_ heterotrimeric G protein complex with the Nb35 nanobody

To elucidate the basis for selective recruitment of Gα_s/q_, we complexed ADGRF1 CTF with a miniGα_s/q_ heterotrimeric G protein complex and the Nb35 nanobody, purified the complex by size-exclusion chromatography (Fig. 2a), and determined its structure by cryo-EM to 3.4 Å resolution (Extended Data Fig. 3–4). The cryo-EM map was of sufficient resolution to permit the building of an atomic model for the receptor, G-protein complex, and nanobody (Fig. 2b, Extended Data Fig. 5). The cryo-EM map in the orthosteric pocket on the extracellular face of the receptor permitted unambiguous modelling of the TA (Extended Data Fig. 5), which adopts a short alpha-helical motif with a loop capping the TA *en route* to TM1, akin to other resolved TA structures of aGPCRs^10–13^. The polypeptide backbone of the 7TM domain of ADGRF1 bound to miniGα_s/q_ has an RMSD of 0.6 Å when aligned to the 7TM domain of ADGRF1 bound to either miniGα_s_ and miniGα_i_, highlighting that the overall organization of these structures is highly similar despite their different G-protein partners. An LPC lipid is present at the intracellular interface of TM helices II, III and IV, as seen in ADGRF1 structures bound to miniGα_s_ or miniGα_i_^13^. The acyl chain is similarly positioned in our structure, whereas the headgroup has the phosphate embedded deeper in the detergent micelle (Fig. 2b, Extended Data Fig. 5).

**Figure 2.**
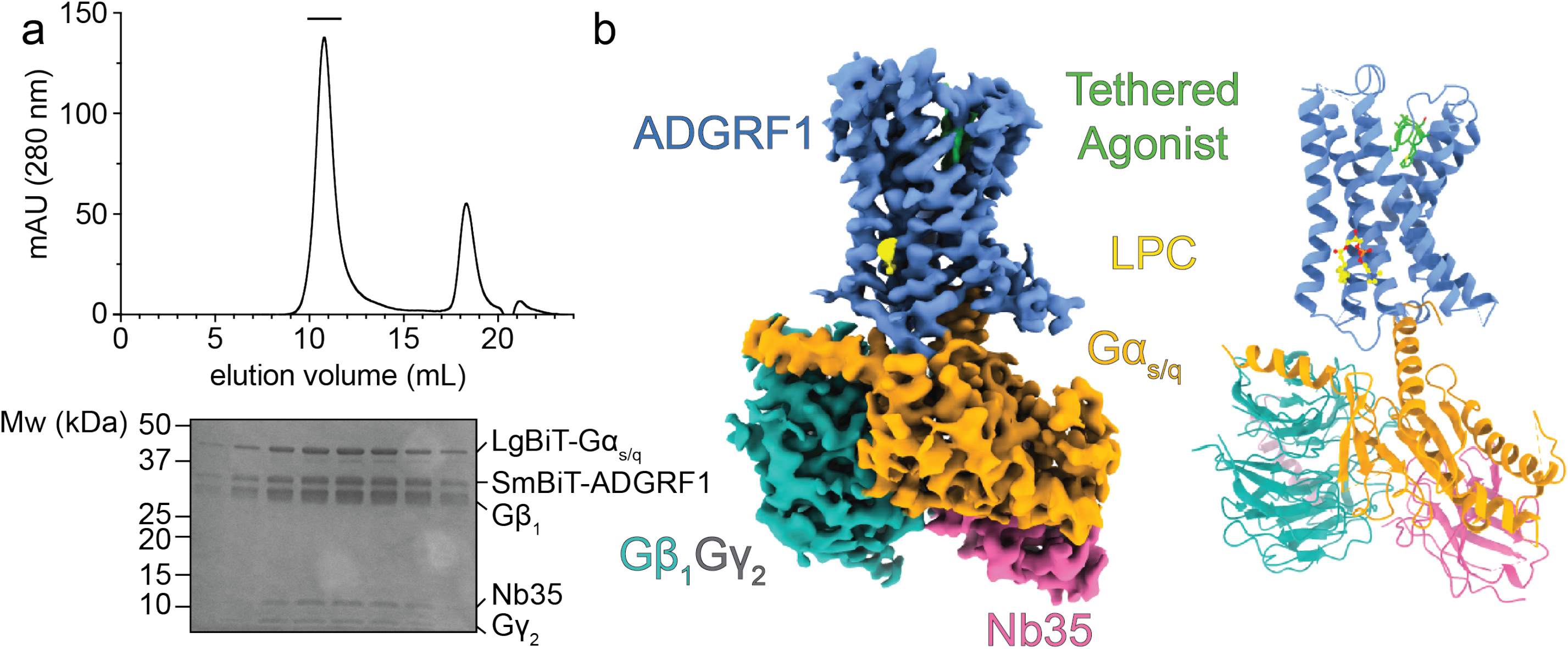
ADGRF1-miniGas/q-Nb35 complex purification and cryo-EM structure. (a) ADGRF1 miniGα_s/q_-Gβ_1_γ_2_-Nb35 purification. Top: elution profile of the complex after size-exclusion chromatography as a final purification step. Bottom: peak fractions (bar above chromatogram) analyzed by SDS-PAGE. (b) Structure of the ADGRF1-Gα_s/q_-β_1_γ_2_-Nb35 complex. Left: cryo-EM map of the ADGRF1 complex. ADGRF1 (light blue), the TA (green), Gα_s/q_ (orange), Gβ_1_ (teal), Gγ_2_ (grey) and Nb35 (pink) are coloured as indicated. The density threshold was set at five standard deviations from the mean. Right: cartoon rendering of the structure, with residue side chains of the tethered agonist rendered as sticks. The density threshold was set at three standard deviations from the mean for the TA. The LPC lipid (yellow) molecule is shown with a ball-and-stick model.

### Structural differences between the ADGRF1 miniGαs/q and ADGRF1 miniGαs complexes

MiniGα_s/q_ was engineered as a chimera with six key Gα_q_ residues on the C-terminal α5 helix that distinguish Gα_q_ coupling over Gα_s_, even though the bulk of the chimera is derived from miniGα_s_^24^. We also observe a preference for miniGα_s/q_ recruitment to the CTF of ADGRF1, consistent with the previously observed preference for Gα_q_ in GTPγS activity assays^8^, arguing that the preference for Gα_q_ is driven largely by the differences in its C-terminal α5 helix. At the ADGRF1 G-protein interface we observe a canonical pose of the C-terminal α_5_ helix at the open cavity at the intracellular opening of TA activated ADGRF1 (Fig. 2b and Fig. 3a). The N387 amide side chain nitrogen of miniGα_s/q_ approaches within hydrogen bonding distance of the backbone carbonyl of ADGRF1 R68 5^3.57^, and the side chain nitrogen of N392 of miniGα_s/q_ approaches within hydrogen bonding distance of the receptor side chain carboxylate of D84 2^8.47^. The analogous residues of miniGα_s_ are H387 and E392, which do not form the same hydrogen bonding arrangement in the Gα_s_ complex. In the Gα_s_ complex, H387 does not make any polar interactions, and instead E392 approaches within hydrogen bonding distance of the backbone amine of S843^8.48^ (Extended Data Fig. 6a). Given the close proximity of E392 to D84 2^8.47^, It is likely that charge repulsion destabilises the miniGα_s_-bound state by comparison to the Gα_q_-bound state. These interactions are associated with a shift in the positioning of the segment linking TM helix VII to helix VIII relative to the Gα subunit, with the Cα of D392 in the α5 helix of miniGα_s_ situated 1.9 Å closer to the Cα of ADGRF1 S843^8.48^ (Fig. 3c). The arrangement seen in the miniGα_s_ complex is also present in the miniGα_i_ structure. Superposition of helices I and VII of miniGα_s/q_ on miniGα_s_ shows a 3.2 Å displacement of the L384 Cα miniGα_s/q_ relative to that of N384 of miniGα_s_ (Extended Data Fig. 6b).

**Figure 3.**
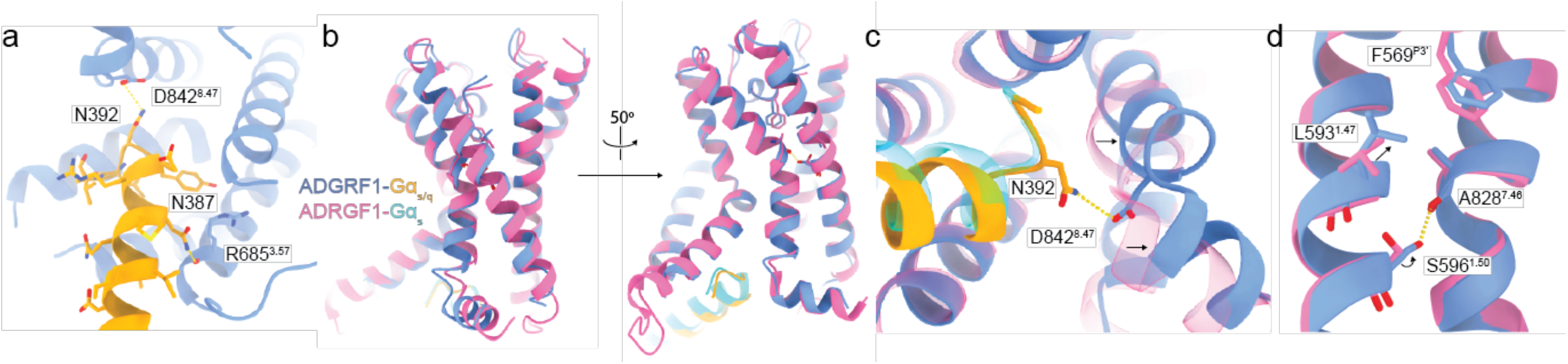
The ADGRF1-miniGαs/q interface and comparison to ADGRF1-miniGαs. (a) Close-up of the ADGRF1 (blue) interface with Gα_s/q_ (orange) along the Cα5 helix of Gα_s/q_. Hydrogen bonding interactions between the receptor and Gα are indicated with yellow dashed lines. (b) Comparison of the ADGRF1 (blue) - Gα_s/q_ (orange) complex with the ADGRF1 (pink) - miniGα_s_ (cyan) complex, aligned by least-squares superposition of the ADGRF1 subunits. (c) Zoomed in view focusing on D84 2^8.47^ in the turn connecting helices VII and VIII. The hydrogen bond between N392 and the carboxylate of D84 2 ^8.47^ is shown with a yellow dotted line. (d) Zoomed in view focusing on residues L59 3^1.47^ and S596^1.50^. Arrows indicate positional shifts of these residues in the miniGα_s/q_ structure relative to their positions in miniGα_s_. The hydrogen bond between S596^1.50^ and the backbone carbonyl of A82 8^7.46^ is shown with a yellow dotted line.

At the orthosteric site, we also observe a 1.2 Å displacement of L.593^1.47^ towards F569^P3’^ in the ADGRF1 complex with miniGα_s/q_, compared with the miniGα_s_ and miniGα_i_ complexes (Fig. 3d). This inward movement leads to a rearrangement of the TM I alpha helix, allowing the conserved S596^1.50^ hydroxyl to approach within hydrogen bond distance of the backbone carbonyl of A828^7.48^ at the kink in the TM VII helix. This interaction of S569^1.50^ with TM helix VII is not observed in either the miniGα_s_ or miniGα_i_ structures.

### Distinguishing preferential Gaq over Gas tethered agonism at the orthosteric site

Because ADGRF1 shows preference for Gα_q_ over other Gα partners^8^, we investigated whether Gα_q_ and Gα_s_ signalling exhibited differential sensitivity to alanine mutants of the receptor stalk residues (Fig. 4a), using NFAT and CRE reporter gene assays for Gα_q_ and Gα_s_ coupling, respectively (Fig. 4b). Positions 574-581, corresponding to P8’-P15’, were largely unaffected by mutation to alanine in NFAT and CRE assays, suggesting this region is not directly important for TA dependent agonism, and therefore functioning mainly as a linker connecting the TA to TM1 in ADGRF1.

**Figure 4.**
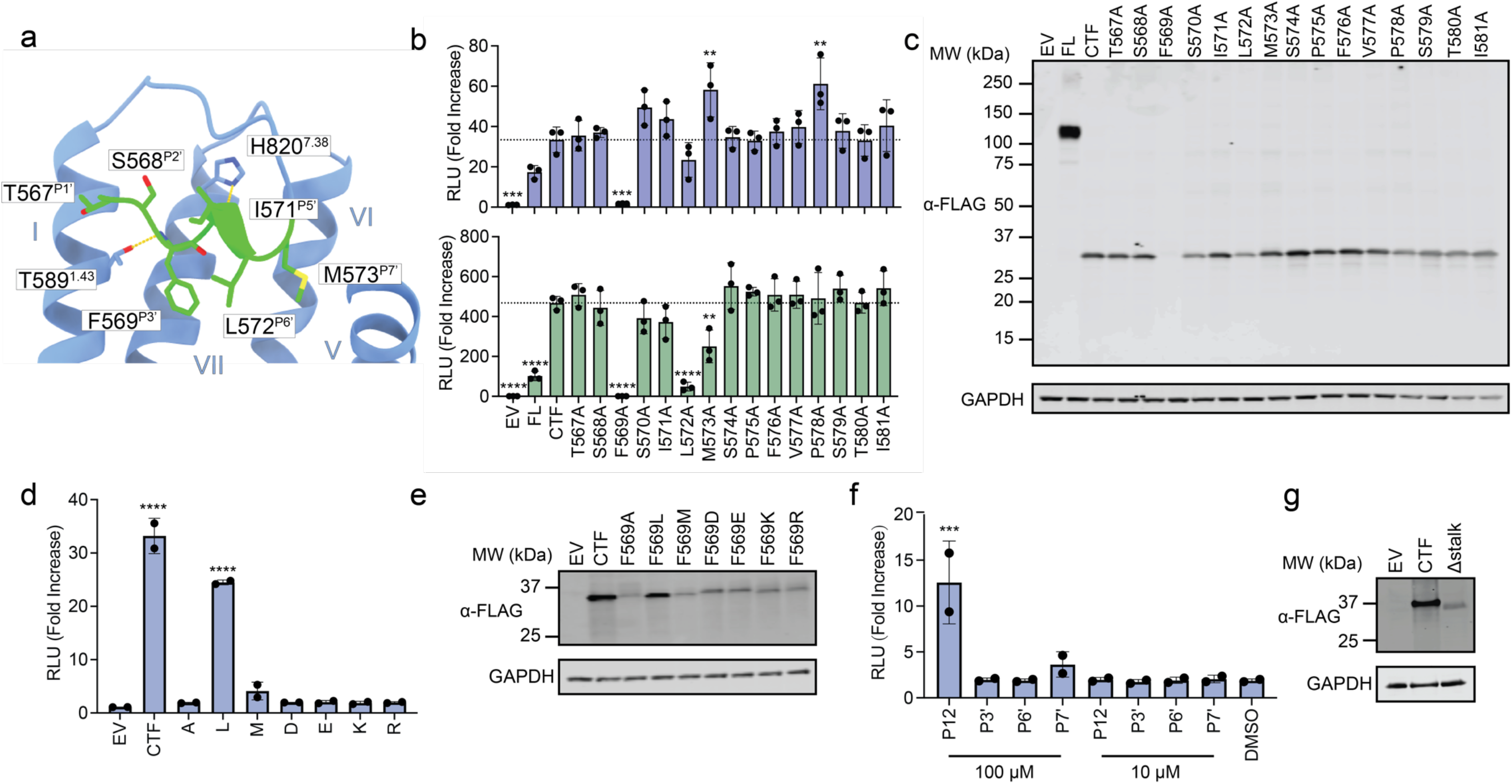
Tethered agonism signalling determinants through NFAT and CRE. (a) Pose of the TA in the orthosteric site. TA sidechain atoms and main-chain atoms making polar contacts are shown as sticks. Side-chains of 7TM residues making polar contacts (yellow dashes) to the TA are also shown. TA superscript labels refer to the TA position relative to the GPS autoproteolysis site. Receptor residue superscript labels refer to Wootten numbering. (b) Scanning alanine mutagenesis of the TA. NFAT and CRE reporter data were acquired using ADGRF1 CTF as a reference sequence. Cells were transfected with 20 ng of receptor DNA/well. Data are normalized to empty vector and reported as relative luminescence units (RLU). The mean for the CTF is indicated as a horizontal dotted line. *Figure legend continued overleaf*. (c) α-FLAG Western blot comparing steady-state amounts of wild-type and mutated ADGRF1 CTF proteins tested in panel b. An α-GAPDH blot was used as a loading control. (d) NFAT signalling data and (e) α-FLAG Western blot of wild-type and P3’ mutated forms (L, M, R, K, D and E) of the ADGRF1 CTF. (f) NFAT reporter gene activity of the ADGRF1 CTFΔstalk protein stimulated by *trans* addition of a 12-residue TA peptide (P12). The response to the wild-type peptide or to peptides with alanine mutations at the P3’, P6’ or P7’ position are shown at doses of 10 or 100 μM. (g) α-FLAG Western blot comparing steady-state amounts of wild-type and Δstalk ADGRF1 CTF proteins tested in panel f. An α-GAPDH blot was used as a loading control. For panel (b), data are presented as mean ± s.d. of three biological replicates. Each point represents the mean value of three technical replicates. For panels (d and f) data are presented as mean ± s.d. of two biological replicates. Each point represents the mean value of three technical replicates. One-way analysis of variance (ANOVA) was used with Dunnett’s multiple-comparison post-hoc test to compare the difference between CTF to all other samples individually in b and d, and to DMSO in f (**P<0.01;***P<0.001;****P<0.0001).

Alanine substitutions at F569^P3’^, L572^P6’^ and M573^P7’^ of the HIM affected TA-dependent signalling. The L572^P6’^ A and M573^P7’^ A mutations both reduced the response of the CRE reporter, whereas these substitutions either did not affect (L572^P6’^A) or modestly increased (M573^P7’^A) the response of the NFAT reporter, arguing that Gα_s_, but not Gα_q_, selectively requires bulky hydrophobic residues at positions P6’ and P7’ for TA-dependent signalling in ADGRF1.

Strikingly, we found that the lack of signalling activity for F569^P3’^ A in both NFAT and CRE is accompanied by greatly decreased receptor abundance, as judged by the lack of a band on Western blot when probing for the N-terminal 3X-FLAG tag with an anti-Flag antibody (Fig. 4c). An antibody raised to a C-terminal portion (residues 831-880) of ADGRF1 showed a similar reduction in Western blot signal for the F569^P3’^A mutant (Extended Data Fig. 7), confirming that the loss of anti-FLAG reactivity was not due to proteolysis of the N-terminal tag. We propose that failure of the mutated TA to engage the 7TM leads to the exposure of the TA hydrophobic sequence, creating a neoepitope that stimulates heat-shock protein binding in the secretory pathway and subsequent degradation (*i.e*., the inactivity of the TA is the cause of the reduced protein amount, and not a consequence of it).

We explored the tolerance of F569^P3’^more extensively by testing a larger range of amino acid substitutions, including a conservative leucine substitution, a methionine substitution, and a suite of charged amino acids (Fig. 4d, e). Among these substitutions, only the F569^P3^’L mutation produced a detectable signal (Fig. 4d), and only this substitution resulted in similar amounts of expressed protein as the wild-type CTF as judged by Western blot (Fig. 4e).

Because of the variation in protein abundance for most of the F569^P3’^ mutants, we also assessed the role of the P3’, P6’, and P7’ hydrophobic buried residues in activation using *trans* addition of TA peptide sequences. For these assays, we used a form of the 7TM domain lacking the TA, which we refer to as stalk-less, or Δstalk. We incubated cells expressing the Δstalk form of ADGRF1 with a 12mer TA mimicking peptide (P12) at 100 μM or 10 μM, and with analogous peptides containing alanine substitutions at P3’, P6’ or P7’, and measured signalling activity using the NFAT reporter. We found that the Δstalk construct signaled robustly through NFAT with the P12 peptide at 100 μM, but not 10 μM (Fig. 4f, g). Together, our data show that a large bulky hydrophobic residue is required at P3’ for ADGRF1 tethered agonism, that Gα_q_ signalling in TA mode is more tolerant of substitutions at the positions at P6’ or P7’ in the HIM motif than Gα_s_ signalling, and that signal activation by peptides added in *trans* is intolerant of alanine substitutions at any of the HIM positions (P3’, P6’, or P7’) even for Gα_q_ signalling.

To further investigate tethered agonism of ADGRF1 through Gα_s_ and Gα_q_, we performed extensive alanine mutagenesis of the TA binding site in the 7TM domain and measured the signalling response of the NFAT and CRE reporters. The signalling activity of the mutated proteins clustered into three distinct categories: mutations that did not affect the CRE or NFAT response, those that were detrimental to both CRE and NFAT, and those that were detrimental to CRE only (Fig. 5a, Extended Data Fig. 8).

**Figure 5.**
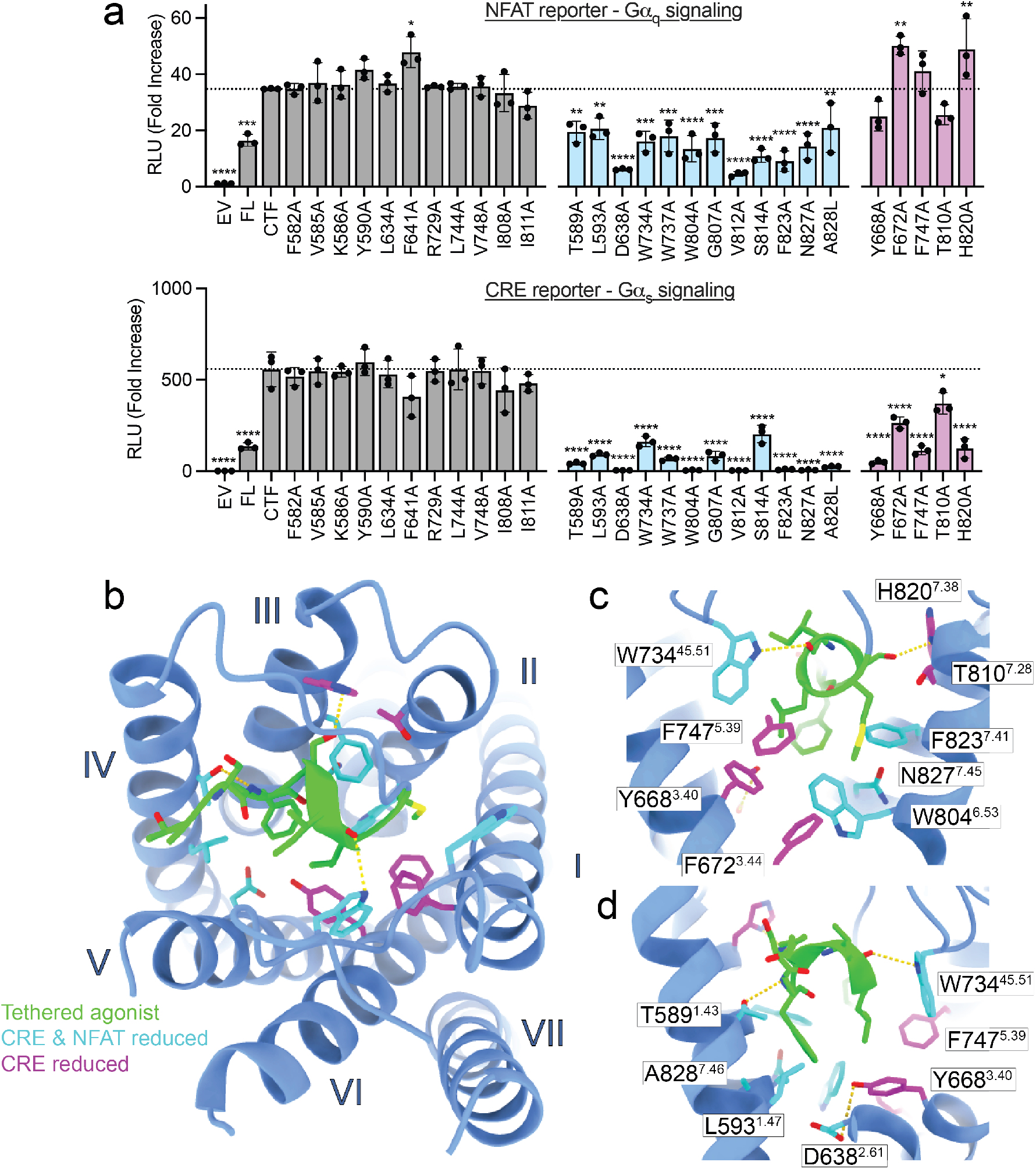
Binding site signalling determinants through NFAT and CRE. (a) NFAT and CRE signalling data for TA binding site mutants in the context of the ADGRF1 CTF protein. Cells were transfected with 20 ng of receptor DNA/well. Data are normalized to empty vector and reported as relative luminescence units (RLU). Data are grouped according to their effects on NFAT and CRE signals relative to the wild-type ADGRF1 CTF protein. Mutations without significant effects on either reporter are grey, mutations that significantly reduce both NFAT and CRE signals are cyan, and mutations that significantly reduce only the CRE signal are magenta. Data are presented as mean ± s.d. of three biological replicates. The mean for the CTF is indicated as a horizontal dotted line. Each point represents the mean value of three technical replicates. One-way analysis of variance (ANOVA) was used with Dunnett’s multiple-comparison post-hoc test to compare the difference between CTF to all other samples individually (*P≤0.05, **P≤0.01, ***P≤0.001 and ****P≤0.0001). *Figure legend continued overleaf*. (b) Top view of the TA in the orthosteric site. TA sidechain atoms and main-chain atoms making polar contacts (yellow dashes) are shown as sticks. Side-chains of 7TM residues that affect signalling when mutated to alanine are shown as sticks and are coloured according to the grouping in (a). (c) View from the P7’ end and (d) the P3’ end of the TA. Side-chains of 7TM residues that affect signalling when mutated to alanine are shown as sticks, coloured according to the grouping in (a), and annotated by residue with Wooten superscripts.

Residues where alanine substitutions reduced signalling in both the NFAT and CRE reporters include conserved residues W73 4^45.51^ and W737 of ECL2, and several residues that surround F569^P3’^ in the 7TM core, including conserved ‘switch’ residue W804^6.53^, neighboring residues F823^7.41^ and N82 7^7.45^, and T589A^1.43^, L593A^1.47^, A828L^7.46^ and D638A^2.61^ (Fig. 5b-d). These sets of mutations outline the importance of ECL2, the binding site arrangement around the ‘switch’ residue, and the conserved F569^P3’^ position for general ADGRF1 tethered agonism. We also found that positions V812^6.61^ and S814^6.63^ lowered signalling in both NFAT and CRE assays, but this was likely due to a profound decrease in protein expression (Extended Data Fig. 8)

Strikingly, mutations that decreased CRE signalling but not NFAT signalling were in two separate clusters that interact with positions other than F569^P3’^ of the TA sequence. The first cluster contains residues within van der Waals contact of L572^P6’^ and M573^P7’^ at positions Y668^3.40^, F67 2^3.44^ and F747^5.39^ on TM helices III and V, with the F672^3.44^ A substitution actually enhancing the NFAT response (Fig 5b-d). The second cluster consists of T810^7.28^ and H82 0^7.38^; T810^7.28^ interacts with M573^P7’^ and S574^P8’^, and H820^7.38^ approaches within H-bonding distance of S570^P4^. These results show that CRE signalling is more sensitive that NFAT signalling upon disruption of the pocket proximal to P6’ and P7’, as well as upon disruption of interactions with S570^P4’^ and S574^P8’^.

## Discussion

For a typical G-protein in which the agonist is a soluble ligand, G-protein coupling in a biological system is guided by intrinsic G-protein binding preferences, agonist selectivity for G-protein, and the local reservoir of G-proteins in that cellular environment. aGPCRs are unique in that the 7TM bundle and its TA are encoded in the same polypeptide chain. As a result, aGPCR G-protein signalling is simplified to a combination of two factors – the G-protein preference of the TA-engaged CTF and the G-protein context of the cellular environment.

Here, we combine cryo-EM structural studies with mutational analysis and cell-based signalling assays to elucidate structure-function relationships in ADGRF1 and deduce its G-protein coupling profile preferences. Our work establishes that ADGRF1 prefers coupling to Gα_q_, as reported previously^8^, provides a structure-based rationale for this coupling preference, and, most strikingly, identifies a cluster of residues in the TA and in the binding pocket that selectively suppress Gα_s_ coupling upon mutation while retaining Gα_q_ coupling (Fig. 6).

**Figure 6.**
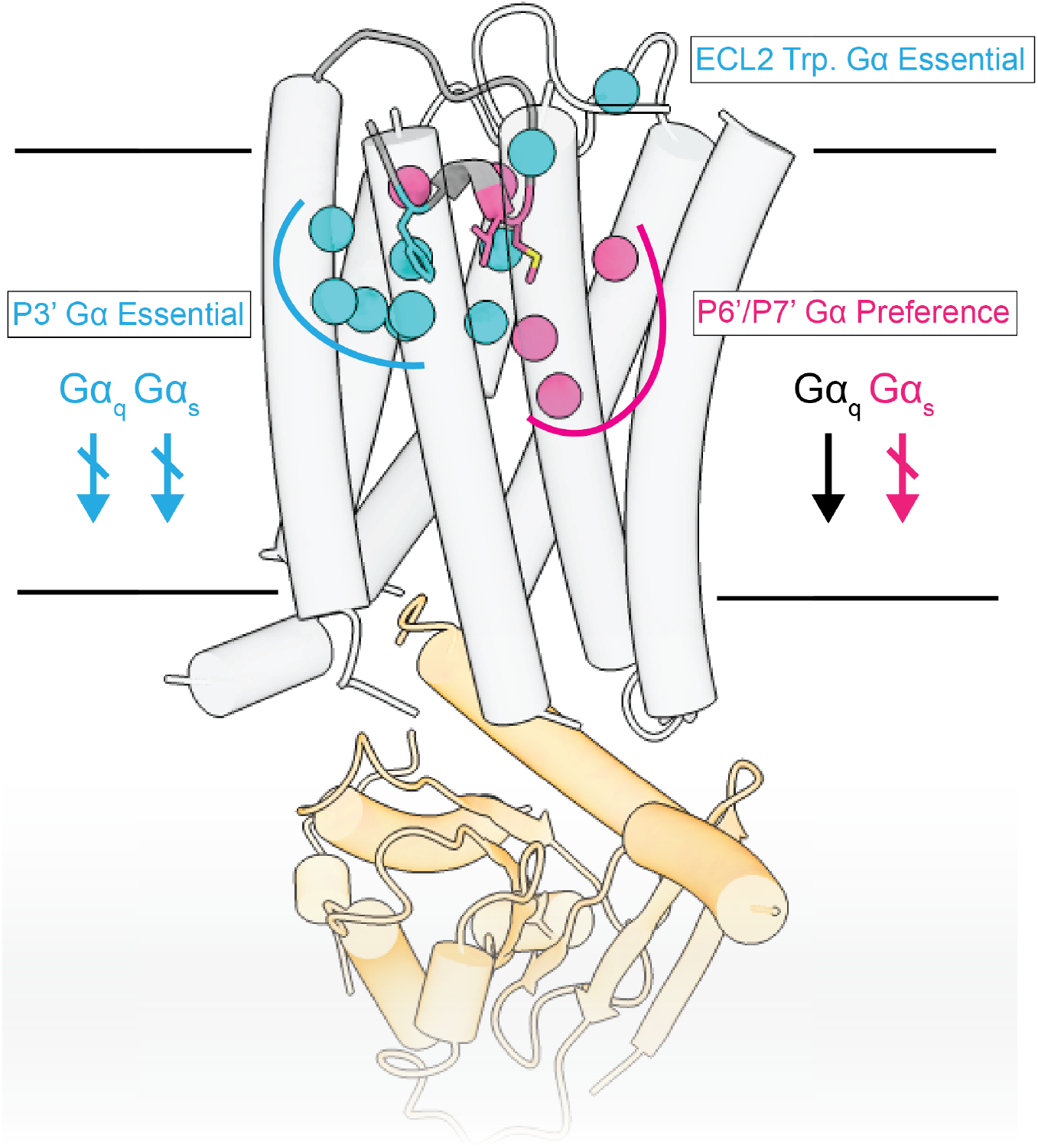
Mutations of the TA and associated binding site cluster by G-protein class preference. ADGRF1 coupled to miniGα_s/q_ represented as tubed helices, colored light grey and orange respectively. Residues that reduced signaling activity are shown, with TA residues rendered as sticks and with Cα atoms of residues in the TA binding site rendered as spheres. Residues that reduce reduce both Gα_s_ and Gaq signalling are colored cyan, whereas residues that only reduce Gα_s_ signaling are magenta.

While the structure of the ADGRF1-miniGα_s/q_ complex shows that the overall architecture of the complex and the pose of the TA in the orthosteric site are similar to recently observed structures of ADGRF1 when complexed to miniGα_s_ and miniGα_i_^13^, there are subtle differences in the complexes that explain why Gα_q_ coupling is more robust than Gα_s_ coupling. We also establish that ADGRF1 can signal through Gα12 or Gα13, which to our knowledge makes ADGRF1 the first class B GPCR to show the functional capacity to signal through all major G-protein classes^28^.

Among the structural differences, tighter packing of L59 3^1.47^ with the highly conserved phenylalanine (P3’) of the HIM appears to be critical for Gα_q_ preference, because the positional shift of L59 3^1.47^ propagates its effects down helix I, introducing a hydrogen bond between S596^1.50^ and A828^7.48^ to stabilize the kink in TM helix VII. Our functional data suggests that these interactions contribute towards an increased stability of the ADGRF1-Gα_q_ complex relative to the Gα_s_ and Gα_i_ complexes. Both the TA and the binding pocket of the 7TM domain show increased mutational lability for Gα_s_ signalling at several residues, whereas Gα_q_ signalling remains unaffected by those same mutations, showing that Gα_q_ signalling is more tolerant to mutations than Gα_s_. Our data also establish the critical importance of the F569^P3’^ residue for ADGRF1 G-protein signalling, because a bulky hydrophobic residue at the P3’ position is essential for expression and TA-dependent activation of the ADGRF1-CTF and mutations of residues that surround the F569^P3’^ pocket are detrimental in both NFAT and CRE assays.

Unlike TA signaling of ADGRF1, which is tolerant of substitutions at positions other than F569^P3’^, we found that *trans* addition of 12mer peptides to a stalkless form of ADGRF1 mimics the TA only when the wild-type sequence is used at the relatively high concentration of 100 μM. Mutations at the P6’ and P7’ positions of the peptides, which are tolerated in TA signaling assays, failed to activate the stalkless ADGRF1 protein. This observation indicates that the development of peptidomimetic agonists for ADGRF1 will require extensive optimization if high potency is to be achieved in a soluble ligand.

Finally, our data elucidating a molecular basis for the preference of ADGRF1 for coupling to Gα_q_ over Gα_s_ identifies key sites in the 7TM that influence G protein coupling preferences. These findings should also inform the study of G protein coupling preferences and signaling biology for other aGPCRs.

## Methods

### Cloning and construct design

#### MBP-CTF and signalling assay plasmids

DNA sequences encoding full length (20-910) and CTF (567-910) forms of ADGRF1 (Uniprot Q5T601) were subcloned into the pFuse-hIgG1-Fc2 vector behind an N-terminal IL2-signal sequence followed by a HA epitope tag, maltose binding protein and TEV protease site^29^. The CTF sequence was also subcloned to create the M-CTF protein with the N-terminal tags and mannose-binding protein omitted. Firefly reporter plasmids (CRE, NFAT, SRE and SRF-RE) were assembled as described^29^. The pRL-TK plasmid (Promega) was used to express *Renilla* luciferase.

#### Receptor and G-protein plasmids

For recruitment assays, sequences encoding full-length (20-910) and CTF (567-910) ADGRF1 were subcloned into pcDNA3.1(+) with an N-terminal haemagglutinin signal sequence followed by a 3XFLAG purification tag and a TEV protease cleavage site. For the CTF construct the TEV protease site P1’ residue encoded a threonine so that the correct residue would occupy T567 in the CTF (e.g., ENLYFQ/TSFSILM…). The C-terminus has a “GGGGSGGGGSSG” linker followed by a SmBiT tag “VTGYRLFEEIL”^30^. For proteins probed by Western blot the C-terminal SmBiT tag was replaced with a HA epitope tag. Mutagenesis was performed with a Q5 site directed mutagenesis kit (New England Biolabs) or using a modified QuikChange™ method^31^.

MiniG protein sequences (miniGαs, miniGαo, miniGαs/q and miniGα12) with capacity to complex with Gβ_1_γ_2_ (e.g. 399 variants^24^) were subcloned into pcDNA3.1(+) with an N-terminal LgBiT fusion followed by a “GGGGSGGGGSSGEF” linker that includes a translated EcoRI site. For structure determination and mutagenesis studies of ADGRF1, the constructs described above for recruitment assays were used.

#### Protein expression, purification and complexation

ADGRF1 and miniGαs/q were co-expressed in Expi293F cells. Cells were seeded at a density of 3.3 x 10^6^ cells/mL in 900 mL of Expi293 expression media. To transfect cells, equal amounts of receptor (250 ug) and miniG (250 ug) DNA were resuspended in 50 mL opti-MEM, FectoPro (Polyplus) transfection reagent (0.5 mL) was resuspended in 50 mL opti-MEM, the DNA and FectoPro solutions were mixed to give a final 1:1 DNA/FectoPro ratio, and the mixture incubated at room temperature for 10 minutes before adding to cells. Approximately 24 hours later, filter-sterilised Valproic acid sodium salt (Sigma-Aldrich) was added to 5 mM, along with 10 mL of 45 % D-(+)-Glucose solution (Sigma-Aldrich). Cells were cultured for a further 24 hours before they were harvested by centrifugation at 4000 *g* for 15 minutes. The pellet was flash frozen and then stored at −80 °C until purification.

Cells were thawed and resuspended in 250 mL of ice-cold hypotonic lysis buffer (10 mM HEPES, pH 7.5, containing Roche protease inhibitor tablets and 0.1 mU/mL apyrase to prevent separation of the GPCR-miniG complex). The pellet was stirred on ice for 30 minutes and lysed by dounce homogenization. The broken cells were pelleted by centrifugation at 50,000 *g* and resuspended in 225 mL solubilisation buffer (10 mM HEPES, pH 7.5 containing 1 mM MgCl_2_, 2 mM CaCl2, and 100 mM NaCl) with 0.1 mU/mL apyrase and 1:100,000 Benzonase (v/v). The resuspended material was dounce homogenized, and DDM-CHS (10:1 pre-mix, Anatrace) was then added to a final concentration of 1 % (w/v). This solution was stirred for 75 min minutes at 4 °C, clarified by centrifugation at 50,000 *g* for 1h, and passed through a glass microfibre filter. The filtrate was then loaded onto M2 anti-FLAG antibody affinity resin by gravity flow. The resin was washed with 30 column volumes of wash buffer (10 mM HEPES, pH 7.5 containing 1 mM MgCl2, 2 mM CaCl2, 100 mM NaCl, and 0.1 % (w/v) LMNG-CHS), and the protein was eluted by the application of wash buffer containing 100 μg/mL of 3XFLAG peptide. The eluted protein was concentrated using a 100 kDa MWCO centrifugal concentrator (Amicon).

Nb35 and human Gβ_1_γ_2_ were purified according to previously published protocols^32^. ADGRF1-miniGαs/q, Nb35 and Gβ_1_γ_2_ were mixed in a 1:1.15:1.15 ratio and incubated overnight at 4 °C with rotation. The protein mixture was centrifuged at 21,000 *g* for 20 minutes at 4 °C, and then spun through a Durapore^®^ PVDF 0.1 μm column (Millipore). The protein mixture was injected onto a Superdex S200 10/300 GL column equilibrated in SEC buffer (10 mM HEPES pH 7.5 containing 1 mM MgCl_2_, 2 mM CaCl_2_, 100 mM NaCl, and 0.005 % (w/v) LMNG-CHS). The peak fraction containing the complex was collected and concentrated using a 100 kDa MWCO centrifugal concentrator to approximately 8 mg/mL.

#### Cryo-EM sample preparation and image acquisition

The complex was vitrified on QUANTIFOIL^®^ holey carbon grids (400-mesh, copper, R1.2/1.3, Electron Microscopy Sciences) using a FEI Vitrobot Mark IV (FEI, Hillsboro). Grids were glow-discharged, and 3 μL of sample loaded to the grid in a chamber at 22 °C and 100 % humidity. Samples were applied at a force of 15 and blotted for 5-7 seconds before plunge-freezing in liquid ethane.

Data were collected using a FEI Titan Krios at 300 kV with a Gatan Quantum Image Filter with K3 Summit direct electron detection camera in counting mode with a total exposure dose of approximately 54 e^-^/Å^-2^. 50 frames per movie were collected at magnification of 105,000x, corresponding to 0.825 Å per pixel. Micrographs were collected at defocus values ranging from - 0.5 to −1.5 um. The processing scheme is summarised in Extended Data Figure 4, in brief, motion corrected and dose-weighted using the RELION motion correction implementation^33^ and contrast transfer function parameters estimated by CTFFIND4^34^. Low quality micrographs were removed using micassess^35^ prior to particle picking in topaz ^36^. ~4.9M particles were initially identified, leading to ~2.9M particles after 2D classification in RELION ^37^. Two independent 3D classifications in RELION were then performed with different initial references, with the best classes from each subsequently combined and duplicates removed to leave ~1.2M particles corresponding to ADGRF1. Following further 3D classification in RELION ~660k particles underwent particle polishing in RELION ^33^ and further 2D classification followed by masked 3D classification with no alignment using a T value of 20 in RELION. 73,903 particles with strong transmembrane density were identified which resulted in a 3.6Å reconstruction from CryoSPARC non-uniform refinement with contrast transfer function refinement ^38^. Subsequent local non-uniform refinement with a mask excluding the micelle improved the reconstruction to 3.44Å.

#### Atomic modelling and model refinement

Model building was carried out in Coot using overlaid non-uniform refinement and local filtered maps generated in CryoSPARC^39^, as well as deepEMhancer^40^ maps. The ADGRF1-miniGα_s_ coordinate file (PDB: 7WU3) was used as an initial model. The miniGα_s_ sequence was changed in Coot to generate miniGα_s/q_^24^. The coordinates were then manually rebuilt using Coot, and refined using ISOLDE^41^, Phenix Real-Space Refine^42^ and Servalcat^43^. The final models were evaluated using MolProbity. Statistics of the map reconstruction and model refinement are presented in Supplementary Table 1. Structural biology applications used in this project (except CryoSPARC) were compiled and configured by SBGrid^44^.

#### Cell Culture

HEK293T and HEKΔ6 cells were cultured in DMEM supplemented with 10% FBS and 1% penicillin-streptomycin at 37°C in a 5% CO_2_ humidified incubator.

#### Dual luciferase reporter assays

HEK293T cells were seeded in DMEM in clear-bottom white 96-well plates pre-coated with poly-D-lysine the day before transfection. At roughly 70 % cell confluency, the medium in each well was replaced with fresh DMEM. Cells were transfected using Lipofectamine™ 2,000 (Invitrogen) at a ratio of 3 μL per 1 μg DNA in Opti-MEM using the manufacturer’s instructions. Each well was co-transfected with 50 ng firefly reporter plasmid, 1 ng pRL-TK plasmid, and the indicated amounts of receptor DNA. Cells were cultured in the transfection mixture for 6 hours before the media was replaced with complete media. 24 hours after transfection, the media was replaced with fresh DMEM, and the cells were incubated for another 8 hours prior to assay readout. For SRF-RE and SRE read-outs the media was replaced with DMEM lacking FBS. Before measuring luminescence, cells were washed once with PBS, 20 μL of 1X Passive Lysis Buffer was added to each well, and the lysates were incubated with shaking for 15 minutes. Cell lysates were serially pipetted, and the assay plate was centrifuged at 400 *g* for 1 minute. Dual-Glo luciferase measurements (Promega) were read on a GloMax luminometer. The ratio of Firefly: Renilla signal was calculated for each well and normalized to the mean of the EV-transfected controls to give Relative Luminescence Units (RLU). Each assay was performed in technical triplicate wells and then repeated in biological triplicate on different days.

For peptide experiments, transfection was performed as above. Peptides were dissolved in DMSO to give 10 mM stocks and diluted to 10x concentration in Opti-MEM immediately before use. The final DMSO concentration was 1 % (v/v) in all wells. Peptides were added *in trans* at the initial media change 6 hours after transfection, and luminescence readings were acquired 24 hours after transfection. Each assay was performed in technical triplicate wells and then repeated in biological duplicate on different days.

For Gα rescue assays, HEKΔ6 cells were seeded as above the day before transfection. Varying amounts of Gα subunit plasmid (0.3 ng of Gα12 and Gα13; 20 ng of Gαs, Gαq, Gα11, and Gα_16_) were co-transfected with 30 ng firefly reporter plasmid and 0.6 ng pRL-TK plasmid (SRF-RE), and total DNA load per well was balanced with pcDNA3.1. Media was changed 6 hours after transfection mixes were added to the wells as in the standard assay format. At 24 hours after transfection, the media was replaced with DMEM without FBS, and the cells were incubated for another 8 hours prior to assay readout. Each assay was performed in technical triplicate wells and then repeated in biological duplicate on different days.

#### G-protein NanoBiT recruitment assays

HEK293T cells were seeded in DMEM in clear-bottom white 96-well plates pre-coated with poly-D-lysine the day before transfection. Cells were transfected using GeneJuice™ (Sigma-Aldrich) at a ratio of 3 μL per 1 μg DNA in Opti-MEM using manufacturer’s instructions. Equal amounts of receptor and miniG DNA (50 ng) were used per well. Cells were incubated for 48 hours and washed with PBS prior to assay readout. Cells were resuspended in opti-MEM and the Nano-Glo^®^ Luciferase Assay was read out on a GloMax luminometer. Each assay was performed in biological triplicate.

**Table 1.**
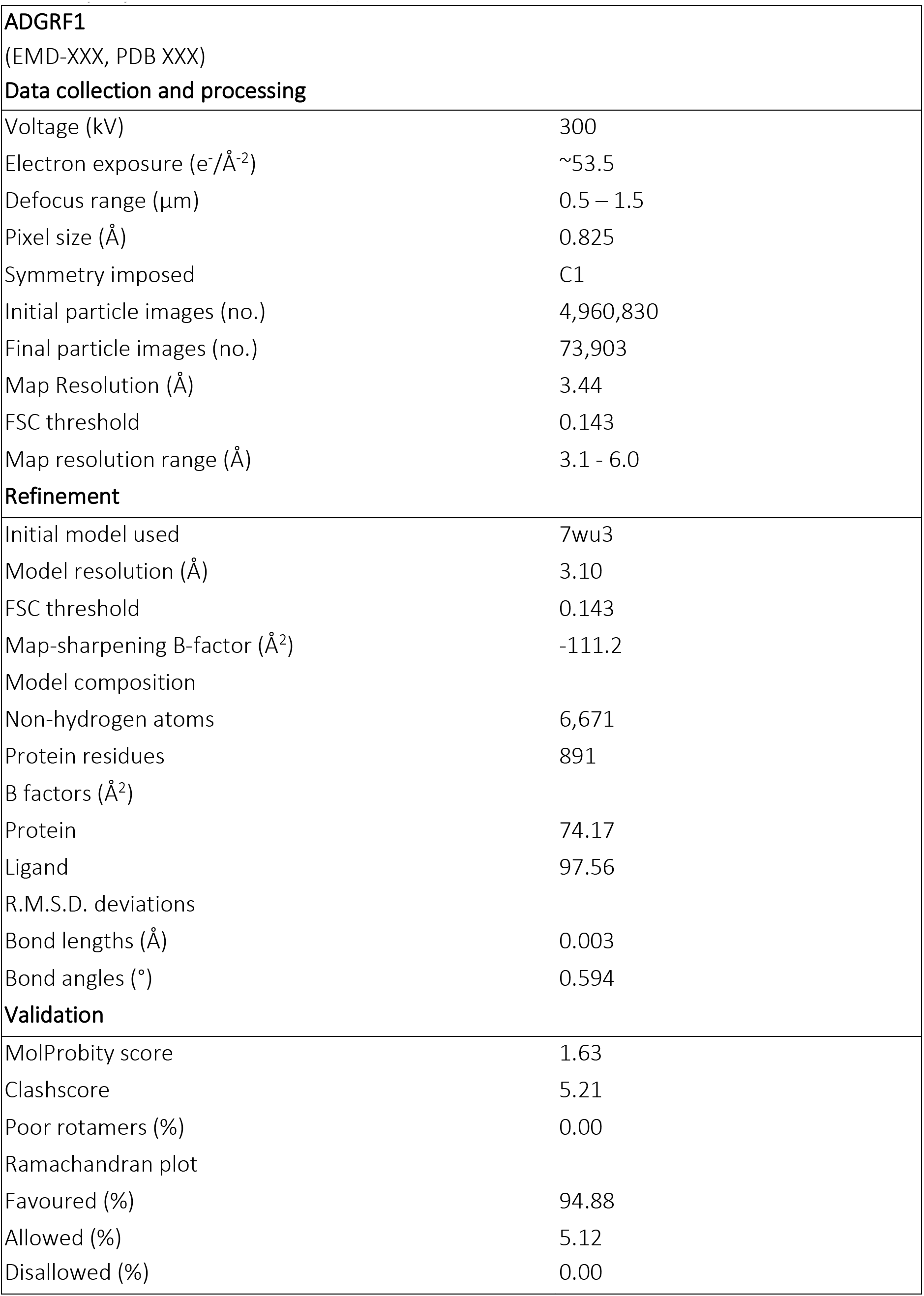
Cryo-EM data collection, refinement and validation statistics.

|  |  |
| --- | --- |
| **ADGRF1** |  |
| (EMD-XXX, PDB XXX) |  |
| **Data collection and processing** |  |
| Voltage (kV) | 300 |
| Electron exposure (e<sup>-</sup>/Å<sup>-2</sup>) | ~53.5 |
| Defocus range (μm) | 0.5 – 1.5 |
| Pixel size (Å) | 0.825 |
| Symmetry imposed | C1 |
| Initial particle images (no.) | 4,960,830 |
| Final particle images (no.) | 73,903 |
| Map Resolution (Å) | 3.44 |
| FSC threshold | 0.143 |
| Map resolution range (Å) | 3.1 - 6.0 |
| **Refinement** |  |
| Initial model used | 7wu3 |
| Model resolution (Å) | 3.10 |
| FSC threshold | 0.143 |
| Map-sharpening B-factor (Å<sup>2</sup>) | -111.2 |
| Model composition |  |
| Non-hydrogen atoms | 6,671 |
| Protein residues | 891 |
| B factors (Å<sup>2</sup>) |  |
| Protein | 74.17 |
| Ligand | 97.56 |
| R.M.S.D. deviations |  |
| Bond lengths (Å) | 0.003 |
| Bond angles (°) | 0.594 |
| **Validation** |  |
| MolProbity score | 1.63 |
| Clashscore | 5.21 |
| Poor rotamers (%) | 0.00 |
| Ramachandran plot |  |
| Favoured (%) | 94.88 |
| Allowed (%) | 5.12 |
| Disallowed (%) | 0.00 |

## Author Contributions

D.T.D.J. and S.C.B. conceived of the research. D.T.D.J., M.M.B. and A.N.D. designed and cloned constructs. M.M.B. performed mutagenesis of TA and binding site residues. M.M.B., A.N.D. and D.T.D.J. performed signaling assays. D.T.D.J. performed NanoBiT recruitment assays. D.T.D.J. expressed, purified and formed the ADGRF1-MiniGα_s/q_-Gβ_1_γ_2_-Nb35 complex. D.T.D.J. and C.H.L. vitrified ADGRF1 complex on grids. S.D.R. and D.T.D.J. processed cryo-EM data, and S.D.R. produced the final map. D.T.D.J. built the ADGRF1 complex model. D.T.D.J. and S.C.B. wrote the manuscript.

## Acknowledgements

We thank members of the Blacklow and Kruse laboratories for helpful discussions. Cryo-EM data were collected at the Harvard Cryo-EM Centre for Structural Biology at Harvard Medical School. We thank Sarah Sterling and Megan Meyer for helpful comments and advice during grid screening, and Megan Meyer for data collection.

## Extended Data

**Extended Data Figure 1.**
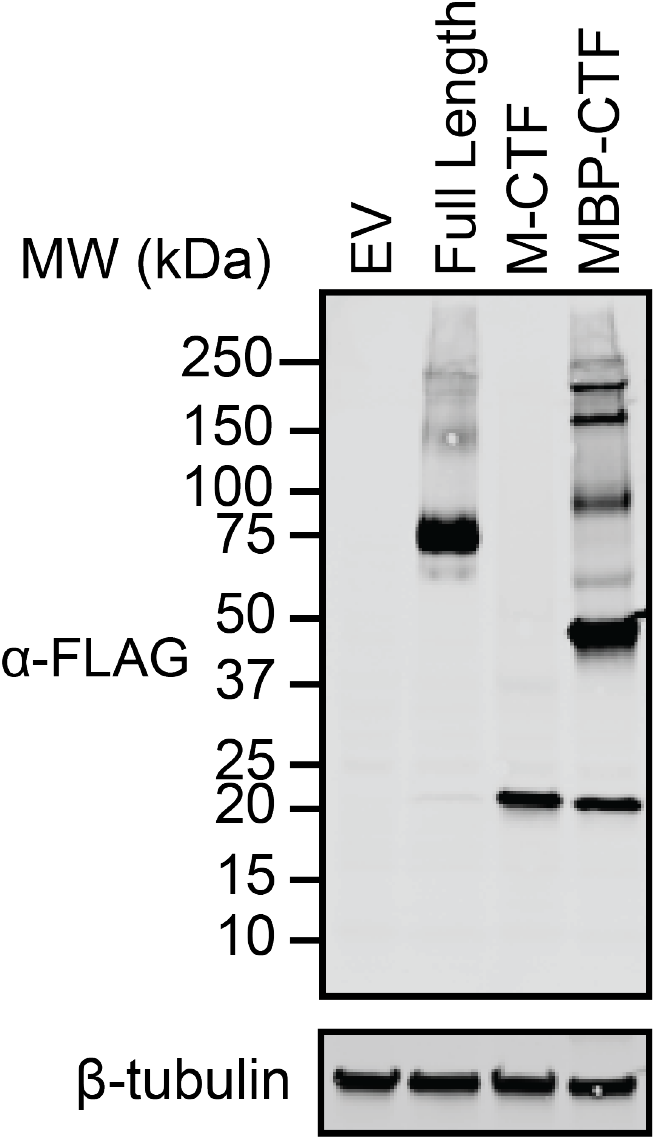
Expression of ADGRF1 proteins used in signalling experiments with NFAT, CRE, SRE and SRF-RE reporters. α-FLAG Western blot of ADGRF1 proteins used in signalling assays to identify G-protein coupling partners. A Western blot with an anti-β-tubulin antibody was used as a loading control.

**Extended Data Figure 2.**
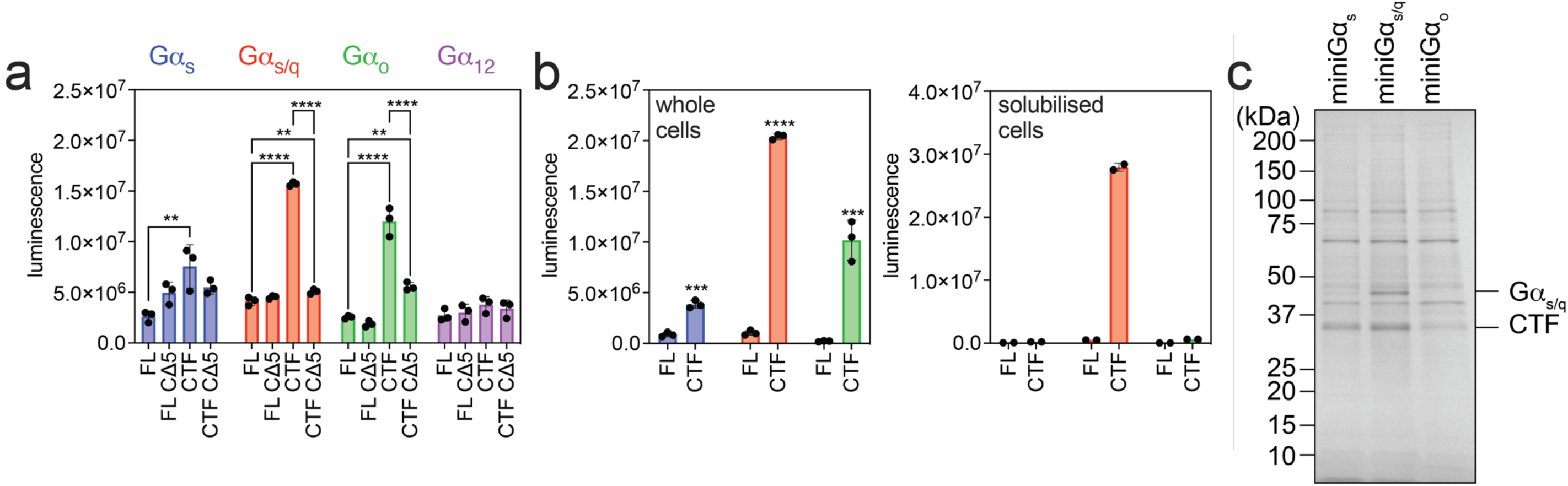
NanoBiT complementation of GPR110-CTF with miniGα proteins. (a) NanoBiT recruitment assays for assessment of receptor-miniG coupling efficiency. Adherent HEK293T cells were transfected with equal amounts of DNA for receptor and miniG proteins and luminescence recorded after the addition of substrate. Full-length (FL) and C-terminal fragment (CTF) variants of the receptor were combined with either functional miniG or non-functional miniG CΔ5 controls. Luminescence resulting from luciferase reconstitution is plotted, with bars grouped and coloured by miniG class. One-way analysis of variance (ANOVA) was used with Dunnett’s multiple-comparison post-hoc test to compare the difference between FL and CΔ5 to all other samples within each miniG group individually (*P≤0.05, **P≤0.01, ***P≤0.001 and ****P≤0.0001). (b) Suspension Expi293T cells were transfected with equal amounts of DNA for receptor and miniG proteins in whole cell (left) and solubilised formats (right). Luminescence resulting from luciferase reconstitution is plotted, with bars coloured by miniG class as in (a). Unpaired two-tailed T-test was used to compare the difference between FL and CTF for whole cell luminescence (***P≤0.001 and ****P≤0.0001). (c) Coomassie blue staining after SDS-PAGE of ADGRF1-CTF complexes recovered on magnetic beads, assessing coupling efficiency to miniG proteins.

**Extended Data Figure 3.**
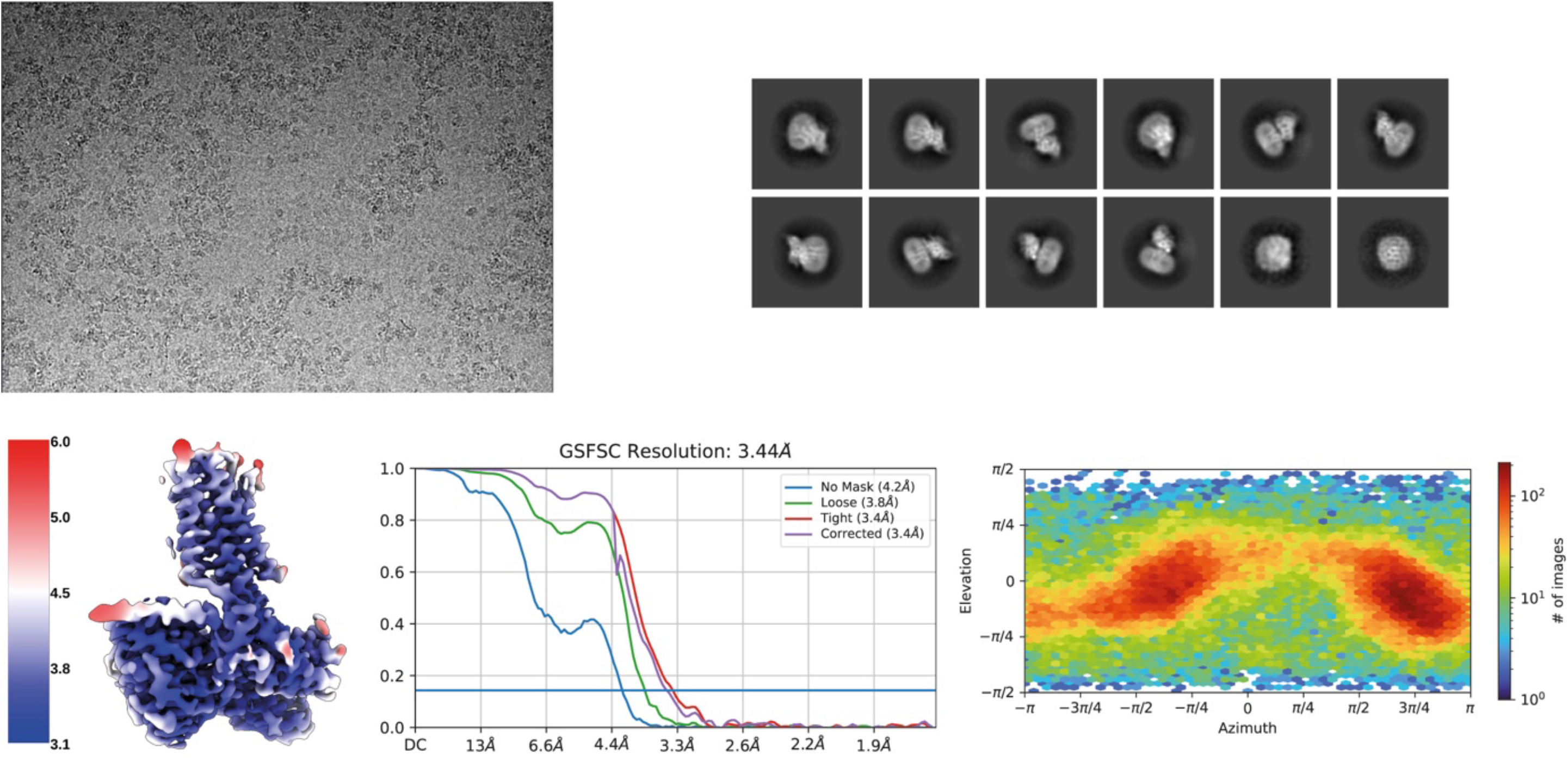
Cryo-EM data quality. (Top, left) A representative image of ADGRF1 complexes in vitreous ice visualized by cryo-EM on a Titan Krios microscope equipped with Gatan K3 detector. (Top, right) 2D-class averages showing a range of particle orientations. (Bottom, left) Local resolution map generated in Cryosparc (GS-FSC = 0.143 cut-off). (Bottom, centre) GS-FSC curves with default Cryosparc masks. (Bottom, right) Particle orientation distribution map of ADGRF1-miniGα_s_/q complexes used for structure determination.

**Extended Data Figure 4.**
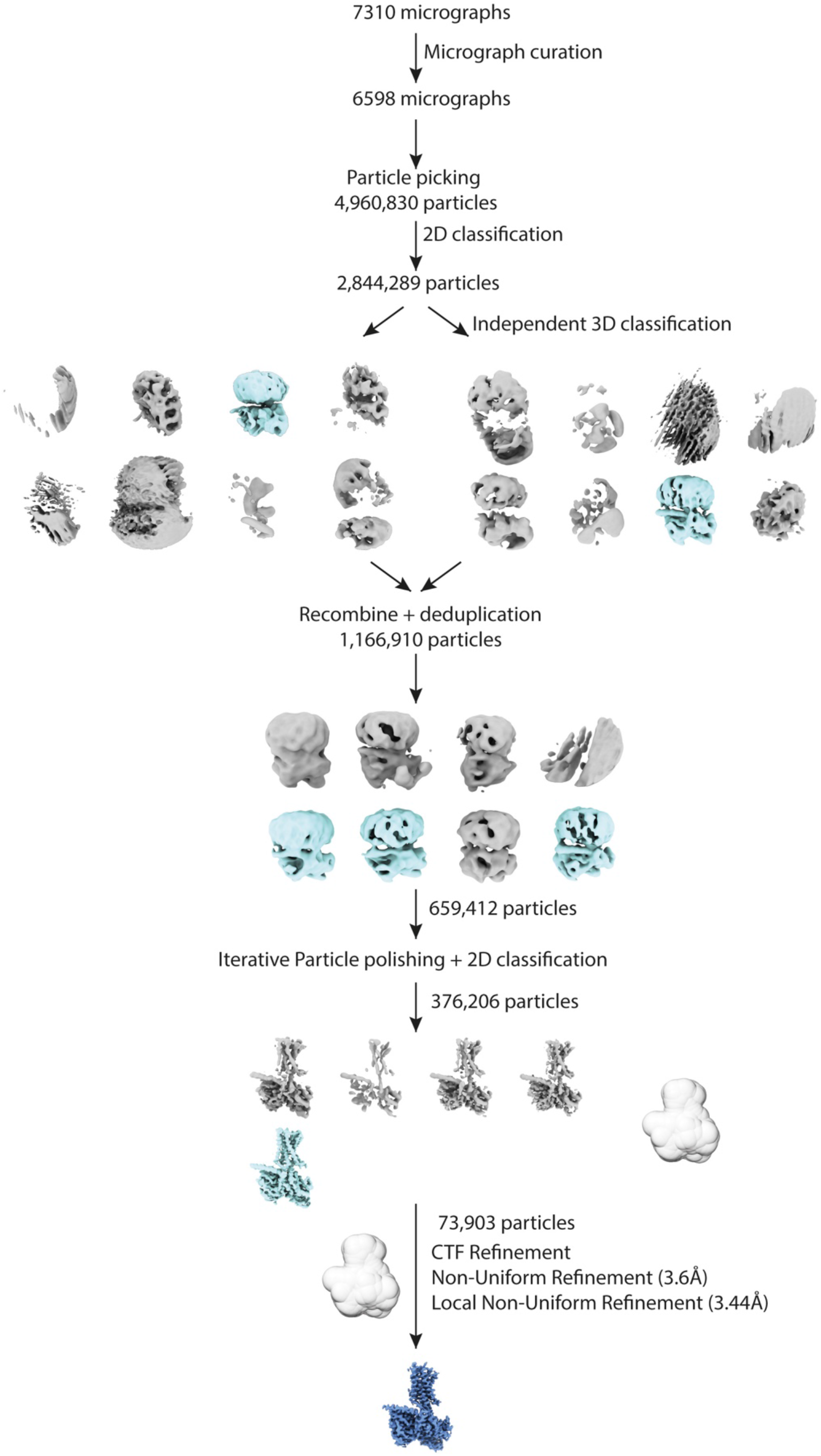
Cryo-EM processing scheme for ADGRF1 reconstruction. Selected classes are blue and discarded classes are grey.

**Extended Data Figure 5.**
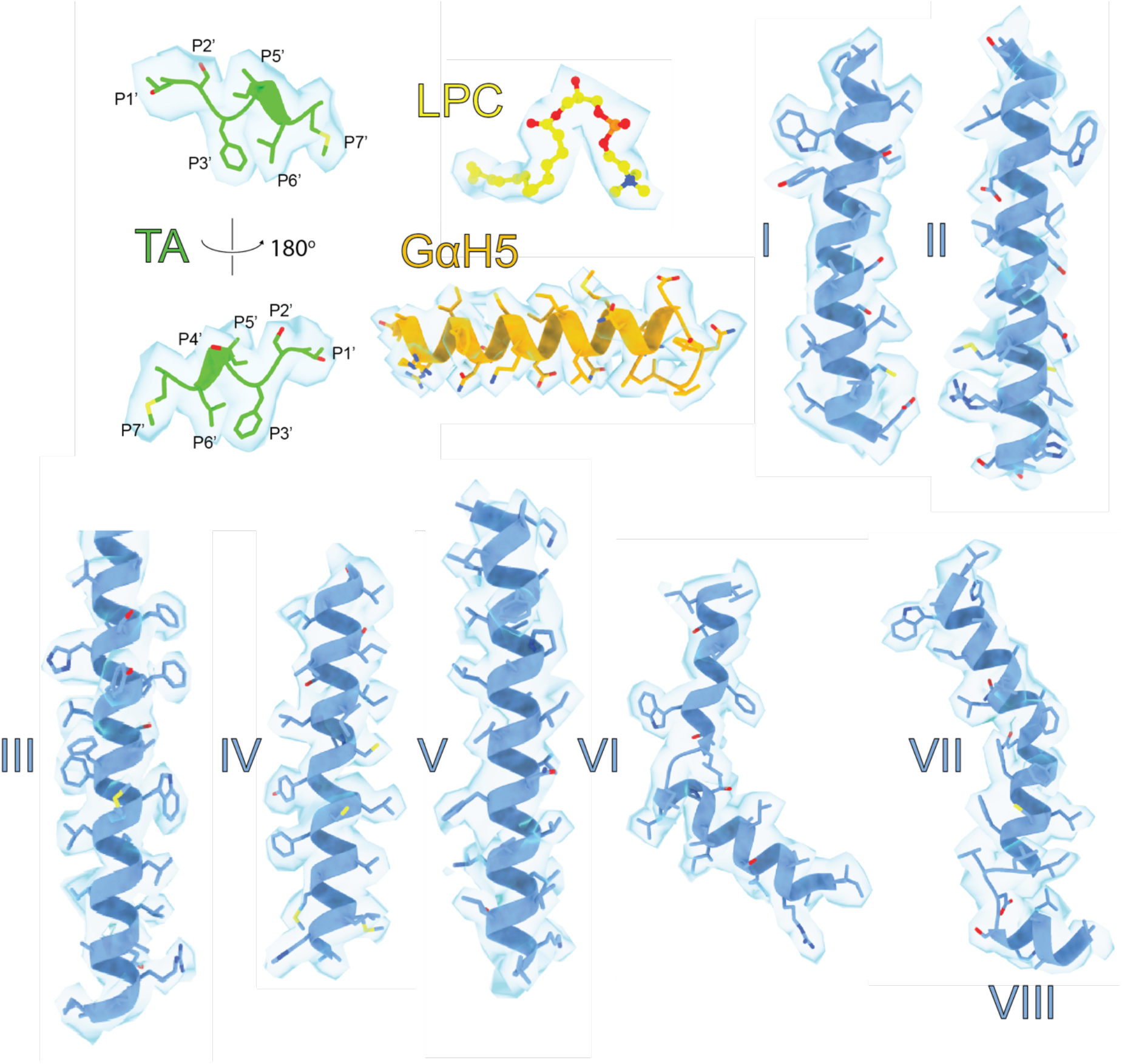
Map to model density of key ADGRF1 components. All protein structures are rendered in cartoon with residue side chains represented as sticks. The LPC molecule is rendered as ball-and-sticks. TA (green); tethered agonist; LPC (yellow), lysophosphatidylcholine (16:0), GαH5 (orange); C-terminal alpha helix of miniGα_s/q_, I-VII (blue) transmembrane helices I-VII, and VIII (blue); helix VIII. The density threshold was set at three standard deviations from the mean for all components.

**Extended Data Figure 6.**
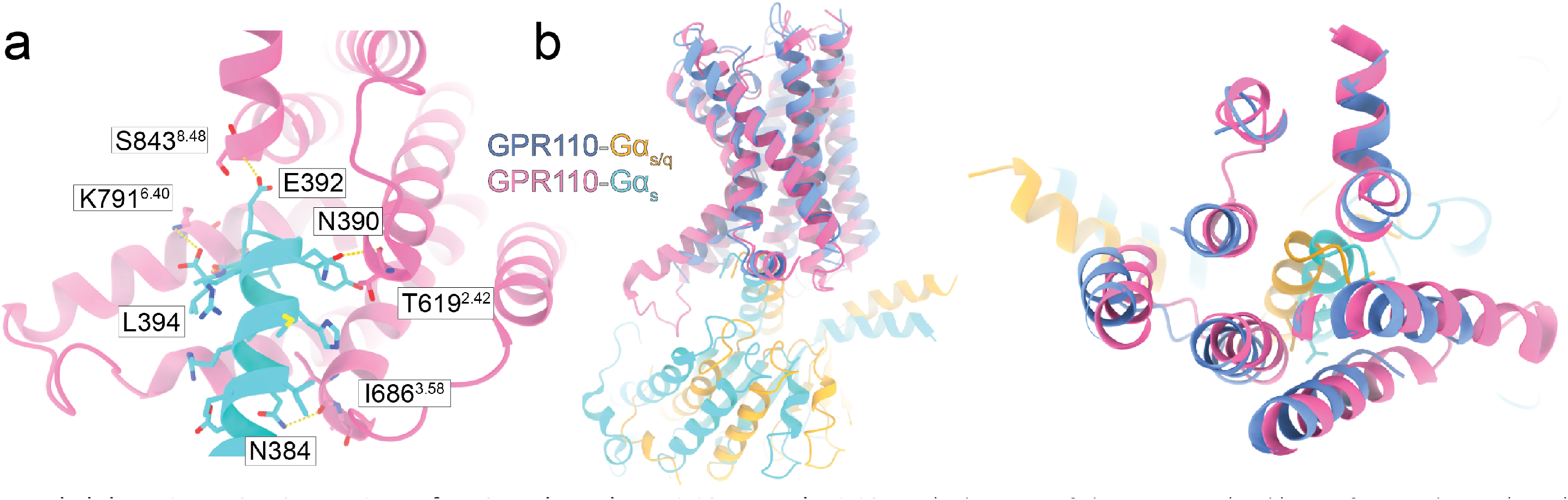
Comparison of ADGRF1 bound to miniGas/q and miniGas. a) Close-up of the ADGRF1 (pink) interface with Gα_s_ (cyan) along the Cα5 helix of Gα_s_. Hydrogen bonds between the receptor and Gα are indicated. (b) Left: side-view, and Right: top-slice view of ADGRF1 bound to miniGα_s_ or miniGα_s/q_. The receptors are aligned by least-squares superposition of TM helices I and VII.

**Extended Data Figure 7.**
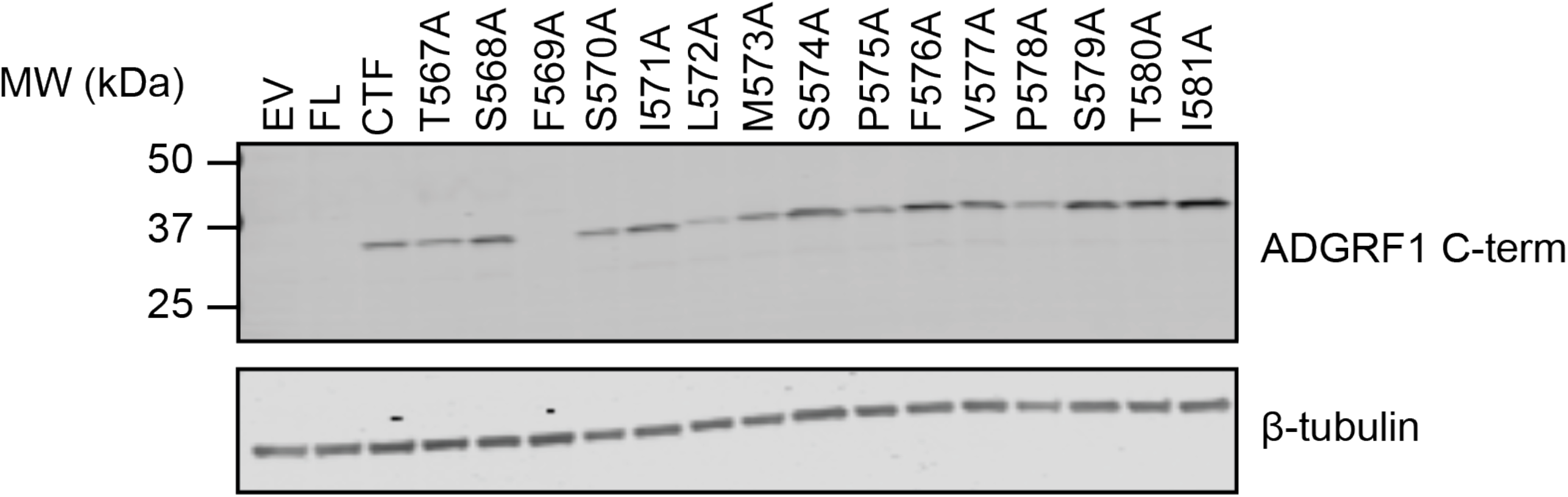
Amount of ADGRF1 CTF protein expressed by wild-type and alanine scanning mutants. Protein was detected by Western blotting with an α-ADGRF1 C-terminal antibody.

**Extended Data Figure 8.**
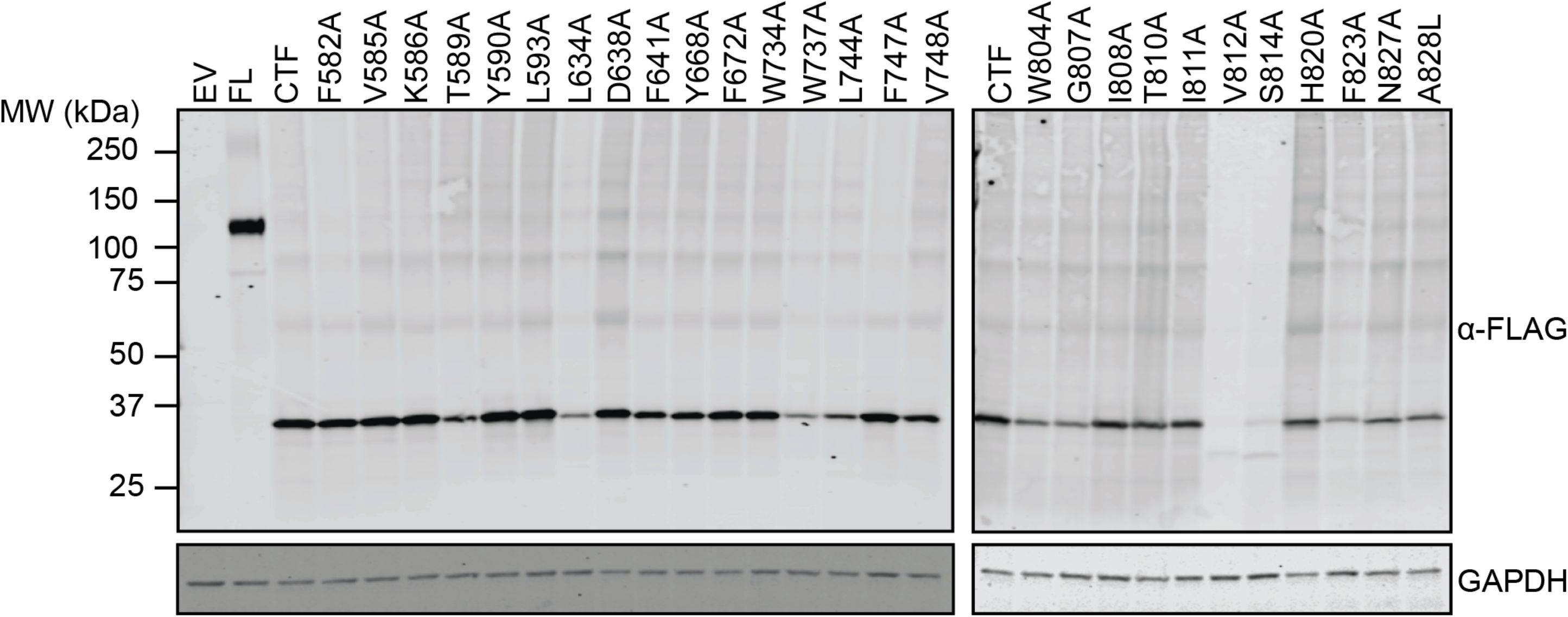
Amount of ADGRF1 CTF protein expressed by wild-type and alanine scanning mutants of the binding site. Protein was detected by Western blotting with an α-FLAG antibody, which recognizes an N-terminal FLAG tag. A Western blot with an anti-GAPDH antibody was used as a loading control. A Western blot with an anti-GAPDH antibody was used as a loading control.

